# Cellular adaptation through fitness-directed transcriptional tuning

**DOI:** 10.1101/137810

**Authors:** Peter L. Freddolino, Jamie Yang, Saeed Tavazoie

## Abstract

Cells adapt to changes in their environment through transcriptional responses that are hard-coded in their regulatory networks. Such dedicated pathways, however, may be inadequate for adaptation to novel or extreme environments. We propose the existence of a fitness optimization mechanism that tunes the global transcriptional output of a genome to match arbitrary external conditions in the absence of dedicated gene-regulatory networks. We provide evidence for the proposed tuning mechanism in the adaptation of *Saccharomyces cerevisiae* to laboratory-engineered environments that are foreign to its native gene-regulatory network. We show that transcriptional tuning operates locally at individual gene promoters and its efficacy is modulated by genetic perturbations to chromatin modification machinery.

## Introduction

The capacity to adapt to changes in the external environment is a defining feature of living systems. Cells can rapidly adapt to familiar changes that are commonly encountered in their native habitat by sensing the parameters of the environment and engaging dedicated regulatory networks to establish adaptive gene expression [1, 2]. However, dedicated sensory and regulatory networks become inadequate, or even detrimental, when cells are exposed to unfamiliar environments that are foreign to their evolutionary history [3]. In principle, at least one gene expression state that maximizes the health/fitness of the cell always exists, despite the inability of the native regulatory network to establish such a state. This is true because under any conceivable environment, the activities of some genes are beneficial, whereas those of others are futile or even actively detrimental [1, 3, 4]. In fact, if the initial fitness defect is not lethal, a population of cells may slowly adapt to an unfamiliar environment through the accumulation of genetic mutations that rewire regulatory networks thereby achieving more optimal gene expression states [3, 5-12].

We wondered whether cells (especially free-living microbes) have evolved alternative strategies for finding adaptive gene expression states on more physiological timescales, without relying on their hard-coded sensory and regulatory systems. In particular, since the perception of the external world may be of limited value under unfamiliar conditions, perhaps a more effective strategy would be to focus on maximizing the internal health of the cell—without regard to the specific parameters of the outside world. This would be a challenging strategy, as every gene in the genome would need to independently discover the expression level that maximizes the internal health of the cell, and these expression levels could vary significantly from condition to condition. In particular, we asked whether individual genes could, in principle, carry out a search process equivalent to gradient descent [13], where the health consequence of stochastic alterations in gene expression could gradually tune the expression of individual genes towards a level that is optimal for internal health. We reasoned that such an optimization process would require the existence of: (**1**) a source of stochastic transitions in gene expression; (**2**) the ability of local chromatin to maintain a record of recent changes in transcription; and (**3**) a central metabolic hub that integrates diverse parameters of intracellular health and continuously broadcasts whether the overall health of the cell is improving or deteriorating. In fact, we find that the foundations for meeting these requirements are already present in eukaryotic cells: (**1**) The expression of many genes is dominated by noisy bursts of transcription—a widespread phenomenon of largely unknown functional significance [14-18]; (**2**) Co-transcriptional histone modification can modify eukaryotic chromatin in promoters and gene bodies, establishing a short-term memory of recent transcriptional events [19, 20]; and (**3**) such global health integrators have evolved in eukaryotes. A classic example is the mTOR pathway, which integrates a diverse array of intracellular parameters including amino acid/nitrogen availability, environmental stress, and ATP levels [21-24].

With the necessary components for gradient-based optimization of gene expression in place (**Fig. 1A**), the promoter of each gene would be able to conduct a simple search process that culminates in finding the expression level that maximizes the overall health of the cell: if global fitness/health is increasing *and* there was a previous increase in transcriptional output (representing larger or more frequent transcriptional bursts), the promoter further increases its transcriptional activity (**Fig 1B**). If fitness is decreasing *and* there was a previous increase in transcriptional activity, the promoter decreases its transcriptional output. Transcriptional output is altered in the opposite direction in the event that there was a previous decrease in transcriptional output. For each gene, this tuning process can be expressed as: *ΔE*_*t*_ = *k*·*sgn*(*ΔF*_*t*_ · *ΔE*_*t–*1_)+ *η* (see **Figs. 1 and 2A**). One can easily see how the process described here can tune the optimal expression of a single gene. What is remarkable, however, is the ability of this hypothetical transcriptional tuning process to find near-optimal gene-expression states for a system with thousands of genes. As can be seen in simulations presented in **Figure 2**, this is achieved through a fitness-directed stochastic search culminating in individual genes finding specific gene expression levels that maximize the health/fitness of the cell. Such a transcriptional tuning mechanism would be highly valuable to free-living organisms, enabling them to optimize their global gene expression patterns to match the specific requirements of any environment in which their dedicated sensory and regulatory networks are inadequate or sub-optimal.

**Figure 1.**
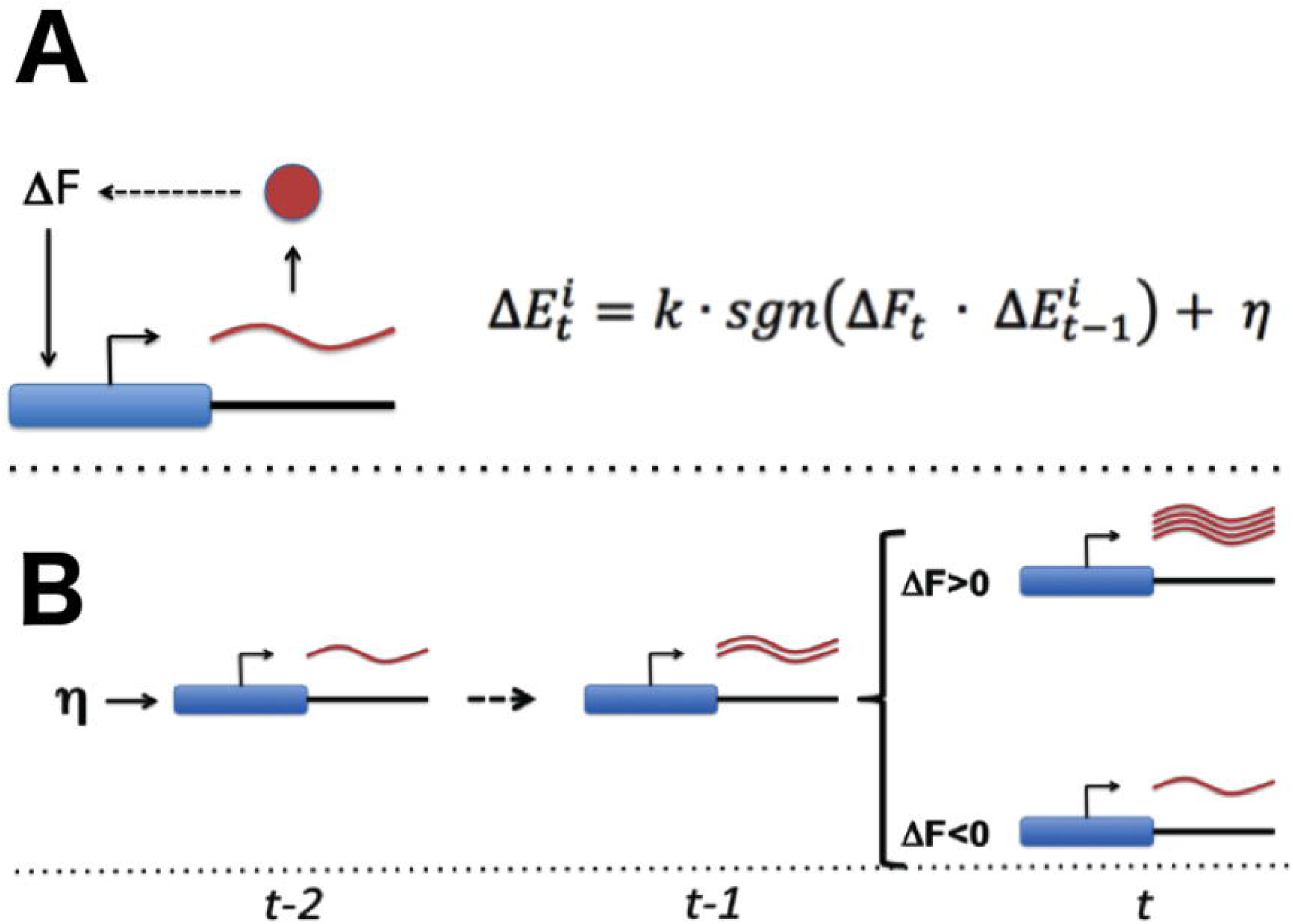
Transcriptional tuning of gene expression by fitness optimization at gene promoters. (**A**) Each gene contains a noisy expression apparatus with noise amplitude of η that allows exploration of a range of transcriptional activities. Each transcription apparatus also maintains a record of its previous change in transcriptional activity (ΔE_t-1_). The change in transcriptional activity has the potential of contributing to a change in global health (ΔF_t_) through the downstream effect of the gene-product’s activity (likely through a multi-step pathway; for example, the biosynthesis of a metabolite that is limiting for growth). A global metabolic integrator can transduce this change in health/fitness to every gene’s expression apparatus. At any point in time, the expression apparatus executes a change in transcriptional activity (ΔE_t_) proportional (*k*) to the sign (*sgn*) of the product of ΔE_t-1_ and ΔF_t_ plus noise (η). (**B**) A simple example of this can be seen for a gene that experiences a random burst in transcriptional activity. If this leads to an increase in fitness the expression apparatus further increases transcriptional activity. Conversely, if there is a decrease in fitness, the expression apparatus decreases transcriptional activity.

**Figure 2.**
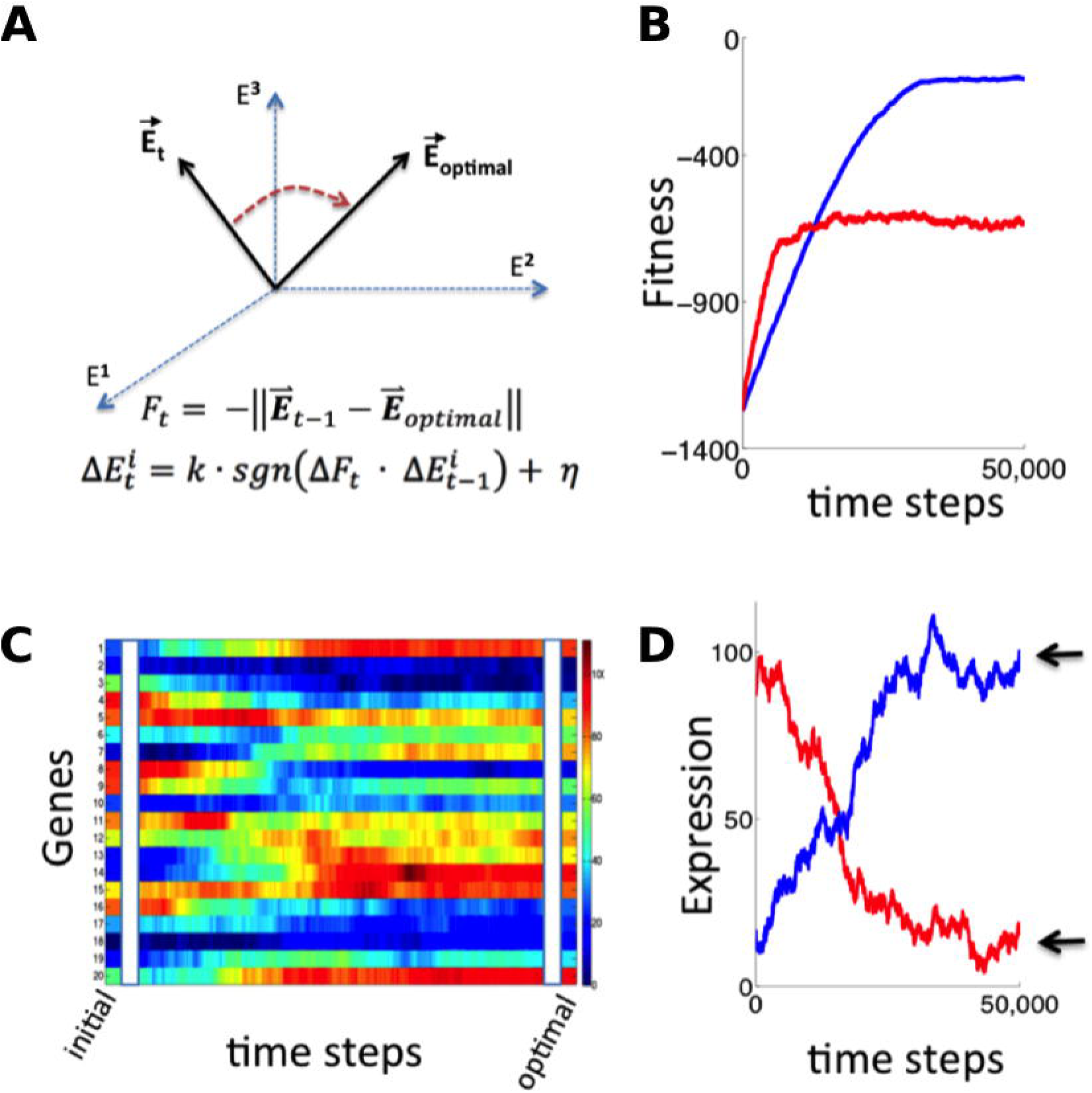
Simulation of fitness-directed transcriptional tuning for a thousand-gene system. **(A)** Quantitative framework describing transcriptional tuning. The transcriptional activity state of the genome is represented by the vector **E,** here schematically represented for a three-gene system. In any environment, there is an optimal transcriptional state vector (**E**_optimal_) that yields maximum fitness. At any time (*t*), a cell with transcriptional activity state **E**_t_ has global health/fitness (F_t_) defined as the negative of the Euclidean distance between the immediately preceding transcriptional activity state **E**_t-1_ and **E**_optimal_. Each gene promoter (*i*) executes a change in transcriptional activity 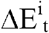 which has two components: (*1*) a step with magnitude of *k* and sign (*sgn*) matching that of the product of the global change in fitness (ΔF_t_) experienced at time *t* and the preceding change in transcriptional activity 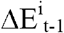, and (*2*) a noise component with a magnitude of η and a random sign (+/-). (**B**) The transcriptional tuning process moves the transcriptional activity state towards the optimum, resulting in increasing health/fitness over time. Simulated trajectories are shown for a 1,000-gene system with *k* =0.1, η = 0.1(blue); *k* =0.5, η =0.5 (red). **(C)** The time evolution of transcriptional activity state vector as a system containing 1,000 genes converges to optimal transcriptional activities through transcriptional tuning. The temporal profiles of 20 representative genes are shown, starting from randomly assigned initial activities, and gradually converging to activities that are near optimal for fitness (using parameters corresponding to the blue curve in panel **B**). (**D**) Trajectories of two representative genes are shown for the same simulation as in panel **C**. Transcriptional activities start at randomly assigned initial values and gradually converge to near the optimum (arrows).

### Fitness-directed tuning of gene expression in yeast

To test for the existence of transcriptional tuning in the eukaryotic model organism *Saccharomyces cerevisiae,* we engineered conditions in which the expression of a single gene was required for growth, but for which no regulatory input existed to drive appropriate expression levels. This was achieved by using a yeast strain (BY4743) which lacks the URA3 gene, which is essential when cells are grown in the absence of uracil. We placed a chromosomally integrated copy of URA3 at a different locus under the control of a weak synthetic promoter, consisting primarily of a pseudorandom sequence. All recognizable binding sites for native transcription factors were removed from the generated promoter sequence (see Materials and Methods and **Data S1** for details), thus decoupling it from any existing sensory and regulatory input. We henceforth refer to this synthetic promoter sequence as synprom. In the experiments described below, URA3 is typically tagged with a fluorescent fusion, either mRuby [25] or a superfolder GFP [26], and a copy of mouse DHFR gene coupled to a different fluorescent protein is inserted at the same location on the sister chromosome to act as an internal control. A schematic of the insertion constructs is shown in **Figure 3A**. We also added the URA3 competitive antagonist 6-azauracil (6AU) to the media to control the threshold level of URA3 production required for growth. The growth condition, SC+glu-ura media, containing *x* μg/ml of 6AU, will henceforth be referred to as ura-/6AU*x.*

**Figure 3.**
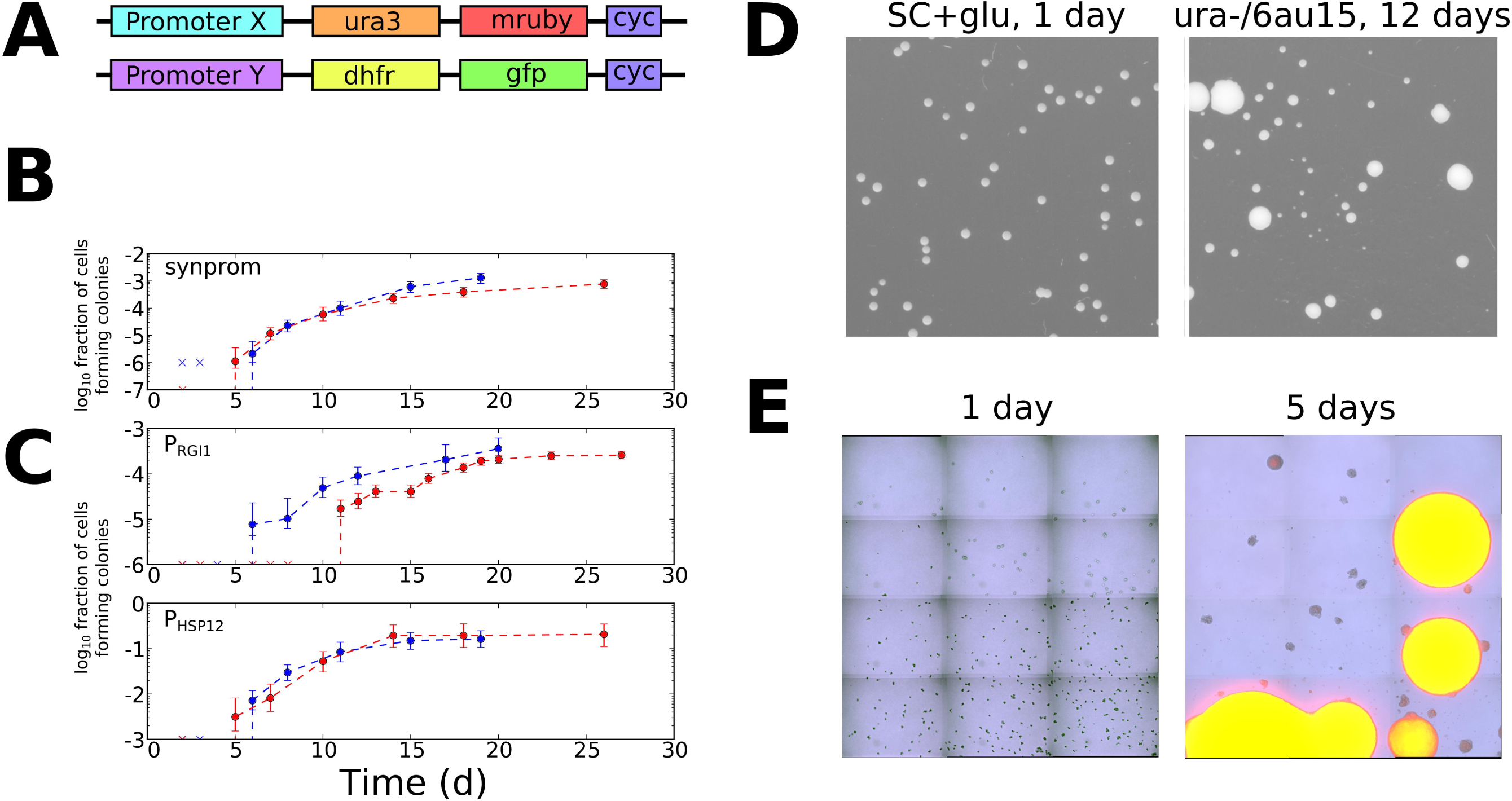
Transcriptional tuning of yeast cells under uracil starvation. (**A**) Schematic of the constructs used in this study. All strains are diploid, containing similar insertions at the LEU2 locus of both copies of chromosome III. X is either synprom or a natural promoter (P_RGI1_ or P_HSP12_) unless otherwise noted, and Y is either the same promoter as X or is the strong constitutive promoter P_ADH1_. “cyc” indicates the well-characterized CYC1 transcriptional terminator [88]. (**B**) Stochastic colony formation on ura-/6AU15 plates for cells containing URA3-mRuby under control of synprom and DHFR-GFP under control of P_ADH1_. Error bars show central 95% credible intervals; colors show biological replicates performed on different days. “x” marks are shown at the bottom of the axis for days where zero visible colonies were present at all plated dilutions. Cells plated on SC+glu uniformly form visible colonies within 1-2 days. (**C**) As in panel B, but with URA3-mRuby controlled by P_RGI1_ or P_HSP12_ as indicated. (**D**) Images of colony growth on SC+glu and ura-/6AU15 plates taken at the specified number of days after plating (1 day for SC+glu, 12 days for ura-/6AU15). Growth of colonies is nearly uniform on SC+glu plates but shows non-uniform stochastic emergence on ura-/6AU15. *N.b.* the plated dilutions for the two plate types are not the same. URA3 expression for the experiment shown is controlled by P_HSP12_, but similar behavior was observed for all promoters discussed here. (**E**) Early colony formation on ura-/6AU15 plates imaged by superimposed differential interference contrast and fluorescence microscopy. Cells contain P_HSP12_-URA3-mRuby/P_ADH1_-DHFR-GFP. Left panel: One day after plating. By this timepoint small, macroscopic colonies would have formed on SC+glu plates, but instead cells remain in microcolonies having undergone no more than three doublings. Right panel: Same plate as left, five days after plating. While most cells have not grown since the one day timepoint, other cells having undergone successful tuning instead form larger colonies with URA3 expression sustained throughout them.

Even with the apparently specific experimental layout described here, with growth highly dependent on URA3 expression, we expect that transcriptional tuning might contribute to fitness through mechanisms acting in *cis* at the promoter driving URA3, those acting in *trans* through modulation of factors that (despite our best efforts) weakly affect the promoter driving URA3, and through tuning of unrelated pathways that benefit survival and growth in the –URA condition. Nevertheless, URA3 expression itself will clearly be the key driver of growth since it is the critical bottleneck for nucleotide biosynthesis in the absence of uracil supplementation.

To look for evidence of fitness-directed transcriptional tuning, we tracked the colony formation of cells containing synprom-driven URA3 after plating on ura-/6AU15 plates. Lacking sufficient URA3 expression to overcome high 6AU levels, these non-growing cells would be expected to succumb to starvation and die. Remarkably, however, after prolonged incubation we observed apparently stochastic transitions to rapid growth, leading to the formation of macroscopic colonies over time (**Figure 3B**). We eventually observed colony formation by roughly one cell in 10^3^, a rate too high to be driven by mutation-driven adaptation in the absence of growth.

### Transcriptional tuning of other synthetic and natural promoters

The synthetic promoter referred to as ‘synprom’ throughout the text is the combination of a pseudorandom sequence with a small natural promoter-proximal region taken from the SAM3 gene, with both stripped of all recognizable matches to known transcription factor binding sites (see Materials and Methods for details). We also tested all combinations of five other synthetic promoter sequences and one other promoter proximal region, enumerated in **Data S2**. As shown in **Fig. S1**, four of the six synthetic promoters support transcriptional tuning, and the ability of synprom5 (the purely artificial component of the synprom referred to in the remainder of the text) itself to undergo tuning remains even with a different promoter proximal region. These findings highlight the universality of the observed tuning mechanism and minimize the possibility that our observations actually arise due to the presence of some residual sequence-specific transcription factor binding site present in synprom.

As shown in **Figure 3C**, we also observed similar tuning behavior for two high-noise natural promoters, P_HSP12_ and P_RGI1_ [27, 28], indicating that transcriptional tuning can function even when superimposed on naturally evolved regulatory sites. Across all promoters (natural and synthetic) tested here, the observed tuning efficiencies varied from 1 in 10^1^ (P_HSP12_) to 1 in 10^5^ (synprom5-arf1).

The apparently stochastic nature of colony formation in our experiments is reflected both in the steady emergence of colonies over the course of days or weeks (**Fig. 3B-C** and **Figs. S1-2**), and in the wide variance of colony sizes observed on ura-/6AU15 plates (**Fig. 3D**). Microscopy revealed that cells remain quiescent for days before transitioning to URA3 expression and rapid growth, with a transition rate dependent on the choice of promoter (**Fig. 3E**). Furthermore, the change thatenables growth under the ura-/6AU15 condition must be passed from mother to daughter cells, as colonies expand from a few points of initiation instead of showing random division of cells throughout the microscopic field over time. While the presence of some deterministic process, yielding colony formation over the observed timescales (dependent on the initial state of each cell), cannot be ruled out, a far simpler explanation for the observed phenomenon of a long lag followed by appearance of colonies over a wide range of times is that each cell independently undergoes a random process that can eventually lead to growth. We confirmed that the appearance of colonies is not simply due to aging of the plates; 6AU-containing plates which were pre-incubated for a week or longer prior to plating of cells showed no change in colony formation rates (data not shown).

### Fitness-directed tuning operates independently of conventional regulatory input and is transcriptionally driven

To provide further insights into the regulatory changes occurring during the onset of cell growth, we performed flow cytometry time courses on cells challenged by, and subsequently growing in, liquid ura-/6AU5 media, using cells with synprom-driven URA3-mRuby, and with a DHFR-GFP fusion driven by either the constitutive ADH1 promoter (**Figure 4A-B**) or synprom (**Figure 4C-D**) itself. The use of P_ADH1_ to drive the second reporter allows us to control for extrinsic noise and global changes in gene expression, whereas coupling synprom to the non-beneficial DHFR-GFP fusion allows us to test whether the observed transcriptional tuning is driven by any *trans-*acting input from some existing regulatory network or whether it is truly specific to the allele needed for growth, as required by our proposed tuning model.

**Figure 4.**
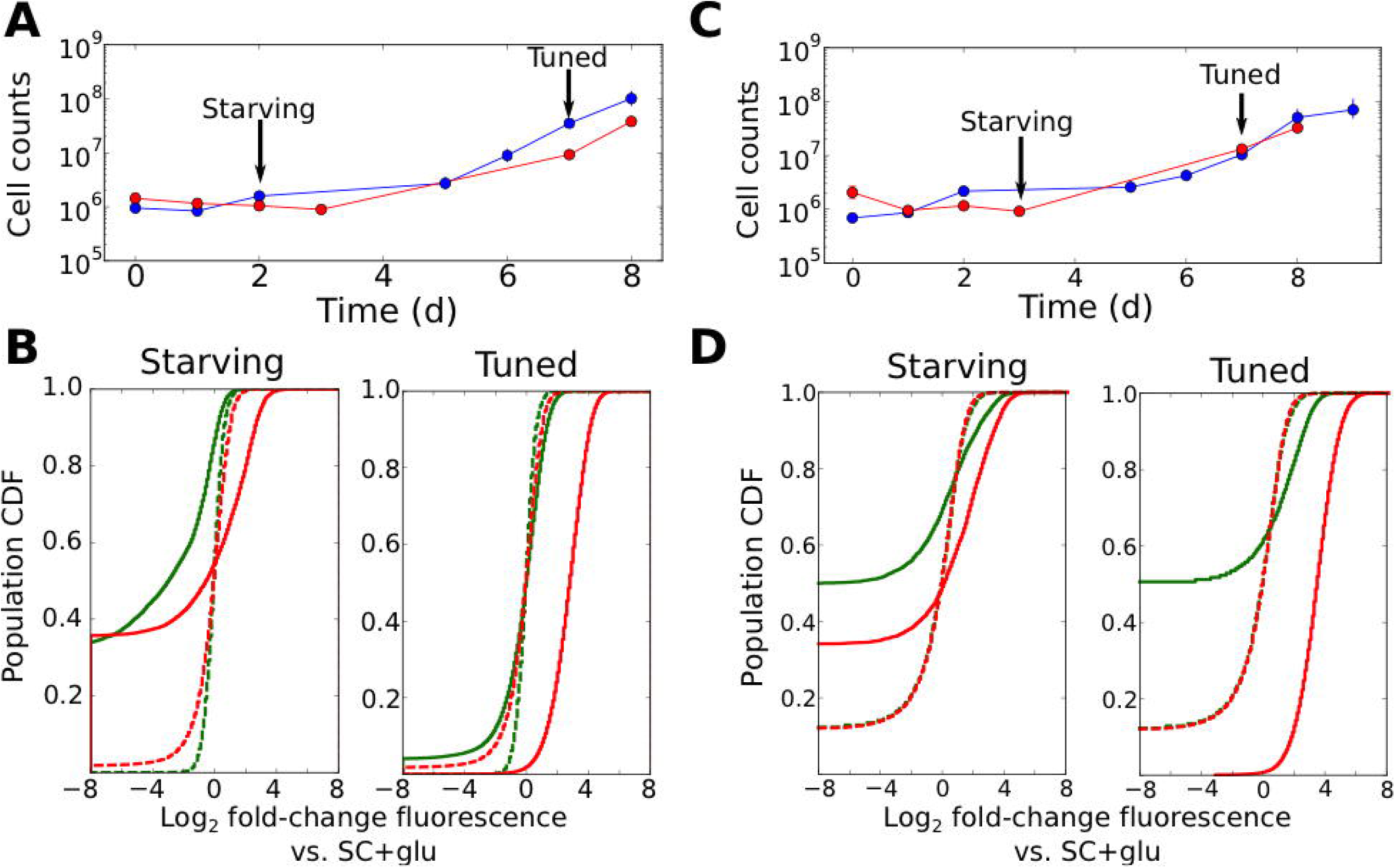
Tuning is both promoter- and allele-specific. (**A**) Cell counts for synprom-URA3-mRuby/P_ADH1_-DHFR-GFP cells in liquid ura-/6AU5 media. Colors correspond to different biological replicates started on different days. Arrows indicate two timepoints from each strain for which fluorescence cumulative distribution functions (CDFs) are shown below. Error bars for cell counts show central 95% credible intervals. (**B**) Flow cytometry cumulative distributions of fluorescence levels for URA3-mRuby and DHFR-GFP during uracil starvation. In each CDF a given timepoint (solid line) is compared to the distribution present for cells in logarithmic growth in SC+glu (rich) media (dashed lines). The values shown are log_2_ ratios to the median value of cells growing exponentially in SC+glu. GFP signals are shown in green and mRuby signals in red. (**C)** Analogous to **A**, but we consider cells where synprom drives both URA3-mRuby and DHFR-GFP. (**D**) Analogous to **B**, but for cells with synprom driving both URA3-mRuby and DHFR-GFP.

Several patterns in the growth curves and flow cytometry data are immediately apparent. First, as with the agar-based growth discussed above, cells show a lag of at least 72 hours with undetectable growth, followed by the onset of steady growth until saturation. With URA3-mRuby driven by synprom and DHFR-GFP by the constitutive promoter P_ADH1_, URA3-mRuby fluorescence increases substantially in tandem with the onset of cell growth, and expression subsequently remains high until saturation; in contrast, DHFR-GFP signals do not even recover to their initial levels (**Figure 4B**; compare dashed and solid line distributions). This demonstrates that the URA3 induction resulting in growth is promoter-specific and does not simply reflect a general increase in protein expression. We observed qualitatively equivalent behavior when URA3 was driven by P_RGI1_ or P_HSP12_ (**Fig. S2**). Even more strikingly, for cells with synprom driving both fluorescent fusions, we observed a specific enhancement of URA3-mRuby expression over that of DHFR-GFP (**Figure 4D**), showing that the transition to high URA3 expression is not only promoter-specific but allele-specific, and thus must be driven at least partly by changes occurring in *cis* at the specific locus whose expression is required for growth. As an additional test, we performed quantitative RT-PCR experiments to measure the ratio of URA3 and DHFR expression in tuned cells either in liquid ura-/6AU5 media or on ura-/6AU15 plates (see **Figure S3**). In both cases, we observed a substantial increase in the URA3:DHFR ratio in the tuned cells, indicating that the observed tuning occurs at least partly through local *cis-*acting mechanisms at the locus required for growth (although we cannot rule out additional changes in other promoters that also contribute to survival and growth, which may account for the observed heterogeneity in expression levels between replicates). Consistent with our proposed tuning model, the allele-specific nature of the transcriptional induction supports a key role for a local tuning process which is independent of dedicated sensory and regulatory input.

### Varying the threshold level of URA3 required for growth shifts tuning from stochastic to deterministic

The presence of the competitive URA3 inhibitor 6-azauracil allows us to vary the threshold level of URA3 required for growth. Thus, it is instructive to consider how the concentration of 6AU may alter transcriptional tuning behavior, both in the context of the computational model described above and in the actual behavior of the system. We made two crucial modifications to the numerical model employed in Fig. 2 to mimic our experimental setup. First, rather than having the entire gene expression profile begin far from the optimal point, we begin with all genes but one (representing URA3) at their optimal values, reflecting the fact that aside from the artificial stress of lacking appropriate URA3 regulation, the cells’ native regulatory network can provide an appropriate response to SC+glu-ura+6AU media. Second, we note that due to the presence of the competitive inhibitor 6AU, the URA3 in the cell will not even be able to contribute meaningfully to uracil production (and thus impact the cell’s health/fitness) until it passes a threshold level. Thus, the tuning term (**Fig. 2A**) is not applied to the gene representing URA3 until after the concentration of URA3 passes a threshold. Aside from the modifications noted above, we model tuning in the ura-/6AU environment as in the general case in **Fig. 2**, and in particular, the fitness effects of changing URA3 expression must compete with noisy gene expression from the other 999 genes in the model gene expression profile to impact the direction of tuning.

The resulting URA3 expression profiles during simulated tuning in the presence of low or high concentrations of 6AU are shown in **Figure 5A**. In the low 6AU case, the tuning mechanism pushes URA3 expression almost deterministically to its optimal (high) value, whereas in the presence of high 6AU, the URA3 expression level undergoes a random walk until expression becomes high enough to allow the tuning mechanism to ‘sense’ the gradient and drive the cells into a URA3+ state. The effects on tuning rates of varying the 6AU concentration are plotted in **Figure 5B**, where we observe that increasing 6AU concentrations both slow tuning and dramatically increase the variance in the amount of time required for each individual cell to reach a URA+ state. This is precisely the behavior observed experimentally with high 6AU concentrations (**Fig. 3**). On the other hand, tuning switched from slow and stochastic to rapid and deterministic in the presence of low 6AU concentrations, with observable tuning occurring over the course of a few hours (**Fig. 5C**). Importantly, the tuning process is confined to the URA3-mRuby allele, despite the fact that DHFR-GFP is also being driven by the same synthetic promoter. This again demonstrates that the tuning process occurs independently of conventional gene regulation by dedicated sensory and regulatory input.

**Figure 5.**
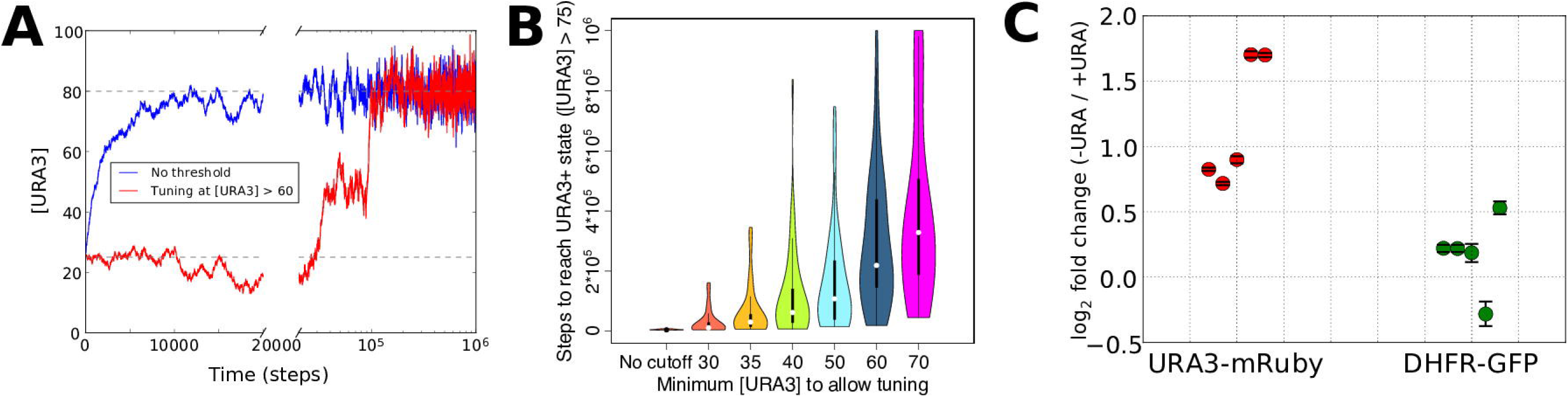
Numerical modeling and experimental validation of changes in tuning behavior as a function of 6AU concentration. We simulated the gene expression dynamics of cells containing URA3 under the control of a non-native promoter, when exposed to uracil-depletion stress with varying concentrations of the URA3 inhibitor 6AU. The model employed is equivalent to that in **Figure 2A**, with *k* = 0.1, η = 0.1, and the target expression profile equal to that for the case shown in **Fig. 2B** except for the case of the gene corresponding to URA3, whose optimal value was set to 80. (**A**) Typical trajectories of URA3 expression levels for a cell in the presence of low (blue) or high (red) 6AU concentrations, which alter the minimum URA3 expression level at which fitness-directed transcriptional tuning can occur. We show results for a starting URA3 level [URA3]=25, with optimal fitness occurring at [URA3]=80. The initial and optimal URA3 levels are shown as gray lines. (**B**) Violin plots of the distributions of the minimum time required to reach a URA3+ state (defined as [URA3]>75) in the presence of increasing concentrations of 6AU (implemented as higher thresholds of URA3 required for transcriptional tuning to become active). In each case profiles reflect 50 independent trajectories simulated at each 6AU level. (**C**) Experimental validation of model predictions. Cells were grown in liquid ura-/6AU1 media (-URA) for 3-4 hours and then had the expression of fluorescent reporter proteins compared (using flow cytometry) with those of the equivalent cells grown in SC+glu (+URA) over the same time period by flow cytometry. Values show log_2_ fold changes from SC+glu to ura-/6AU1; error bars show bootstrap-based 95% confidence intervals. Biological replicates performed on different days are shown side by side; the order of replicates is matched for URA3-mRuby and DHFR-GFP.

### Growth on –ura/6AU media does not arise from genetic mutations

It is crucial to exclude the possibility that genetic mutations underlie the observed tuning transition on –ura/6AU plates. The ongoing emergence of the tuned state in non-growing cells, over the course of many days, makes mutational mechanisms unlikely. In addition, as seen by microscopy (**Figure 3E**), no more than 1-3 cell divisions occur prior to the onset of sustained growth in a small fraction of cells.

Nevertheless, given the phenomenon of stress-induced mutagenesis in non-growing bacterial cells [29], we wished to conclusively exclude any possibility of mutational mechanisms. To this end, we note that changes in URA3 expression occurring due to mutations should be stably heritable in the progeny of the tuned cells, which would not be expected to revert to a URA3 low state even after restoration of uracil in the media. To test the reversibility of the URA3 high state, we designed an experimental setup in which tuned colonies isolated from ura-/6AU plates were grown for varying numbers of passages in uracil-replete media (SC+glu including uracil) and then re-exposed to uracil starvation (**Figure S4**). If any genetic mutation were responsible for increasing URA3 expression in the tuned cells, the phenotype should be stable for many generations. On the other hand, if fitness-directed transcriptional tuning were responsible, cells should revert to a naïve state following sufficient growth in uracil-containing conditions, as they no longer benefit from URA3 expression. As seen in **Figure 6A**, cells with synprom-driven URA3 show reversion toward the naïve colony formation rates upon growth in SC+glu+ura media, with recovery apparent even after a single round of growth on an SC+glu+ura plate, and subsequently becoming stronger with additional SC+glu+ura passages.

**Figure 6.**
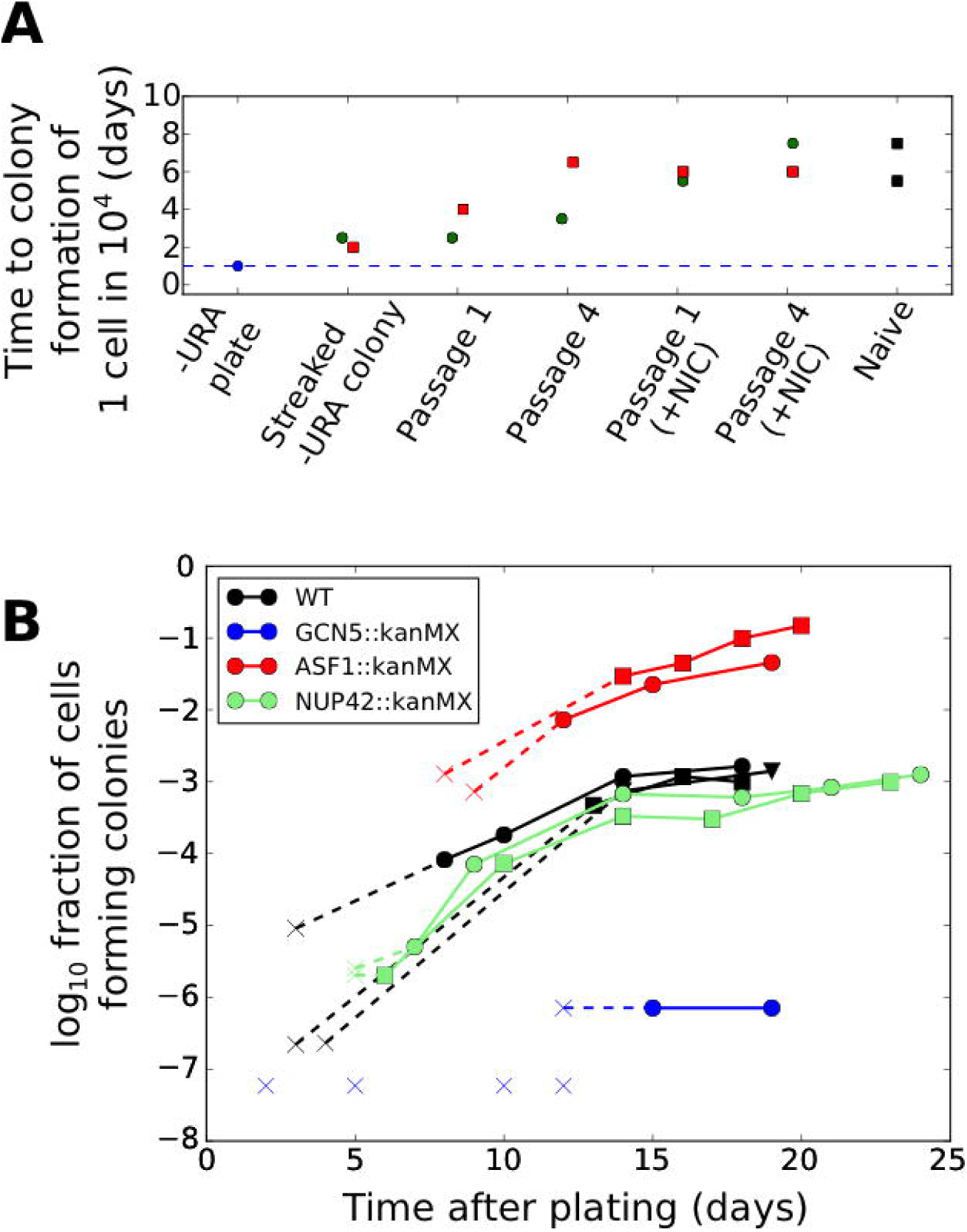
Effects of genetic and chemical perturbations on the efficacy of fitness-directed transcriptional tuning and its epigenetic reversion. **(A)** Time courses of recovery back to the naïve state for tuned cells grown in either SC+glu or SC+glu with 25 mM nicotinamide added (+NIC). Extremes are shown for the colony formation times of cells never exposed to –ura conditions (Naïve) and for single colonies isolated after streaking out cells from ura-/6AU15 plates onto SC+glu (Streaked –URA colony). Colors of points indicate a single lineage beginning from a single streaked out colony picked at the first SC+glu plate stage. The cells were then repeatedly passaged in liquid SC+glu media and assessed for colony formation rates on ura-/6AU15 plates on subsequent days, as detailed in **Figure S4**. **(B)** Colony formation rates on ura-/6AU15 plates in the presence of various genetic perturbations, assessed by colony counts from platings of selected dilutions of cells. An ‘x’ followed by a dashed line indicates no observed colonies, and is shown at the threshold of detection from the experiment. All mutations are in a synprom-URA3-mRuby / leu2Δ0 background.

To conclusively exclude mutational mechanisms, we performed untargeted whole-genome resequencing of a total of eight isolates with synprom-driven URA3-mRuby (four colonies from 6AU15 plates and four separate biological replicates taken after the onset of growth in 6AU5 liquid media; see Materials and Methods for details). For each case, we scanned the region within 25kb of the LEU2 locus (where the URA3 cassettes were integrated) for mutations, since control of URA3 expression was shown in these cells to operate locally in *cis* (**Fig. 4** and **Fig. S3**). The results are summarized in **Table S1**: Of the eight isolates, five show no mutations within 25 kb of the URA3-mRuby insertion, two show SNPs of unknown fitness contribution in a minority of the population, and one shows a duplication of the URA3-mRuby cassette (based on the presence of a read density that is twice the level observed elsewhere for the same chromosome). These data clearly indicate that the origin of growth-supporting URA3 expression levels in these cells cannot be reliant on a mutational mechanism, as only one of the eight cases – that with the URA3 duplication — shows a mutation at high enough levels in the population to explain the onset of growth (mutations present in less than half of the population must have arisen after one or more cells in the population had already tuned and began growing, and thus by definition could not be responsible for the initial onset of the growing state). The phenotypes caused by the sequence variants observed in populations C2 and L4 are not immediately obvious, but even if they are beneficial, their presence in a minority of cells excludes the possibility that they were responsible for the onset of tuning. Note that it should not be surprising (and, indeed, would be expected) that beneficial mutations might arise in a population once it had begun expanding in a new environment due to transcriptional tuning. Our findings are consistent with a non-genetically heritable basis for the observed tuning in seven out of eight of the cases examined.

### Excluding growth-selection on the basis of pre-existing variation in URA3 expression level

A formal possibility for colony formation in a subset of the population is that growth occurs solely on the basis of pre-existing URA3 levels in cells prior to being exposed to uracil deprivation. Microscopic observations of starving cells (**Figure 3E**) argues against this possibility, as a substantial lag passes before any cells begin sustained growth. Also, colony formation continues over the course of many days (**Figure 3D**), demonstrating that even cells that were non-growing for a substantial time period after exposure to URA-stress can eventually grow under this condition. Nevertheless, to conclusively discount the possibility of pre-existing URA3 levels determining tuning, we sorted populations of cells on the basis of initial URA3 expression, isolated those with the highest mRuby levels (the top 0.5-1%, well outside of the main distribution of the population) and plated them. These experiments clearly showed that the ability to form colonies on ura-/6AU plates is not restricted to cells with initially high URA3-mRuby expression (**Table S2**), as the highly fluorescent cells do not form colonies on ura-/6AU plates at rates substantially higher than unsorted cells, and certainly not at a sufficiently higher rate to fully explain the observed colony formation rates. These data argue against the possibility that growth occurs only in cells that, by chance, already have high levels of URA3 expression at the time of plating (although such cells may have some slight advantage, given the nature of their initial state).

### Transcriptional tuning is affected by genetic perturbations to chromatin modification machinery

The proposed fitness-directed tuning mechanism relies on the capacity of local chromatin to maintain a memory of recent changes in transcription and also to modulate the transcription rate based on the fitness consequences of those changes, as conveyed by the proposed central metabolic integrator of internal health. We hypothesized that chromatin modification machinery may be intimately involved in these processes.

To probe the mechanistic basis of transcriptional tuning, we focused on perturbations to histone acetylation/deacetylation (deletions of GCN5, SIN3, HST3, HST4), and chromatin remodeling (deletions of ASF1, ISW2, SWR1, UBP8), all of which provide potential pathways for coupling feedback from the cell’s physiological state to allele-specific modulation of chromatin and transcription (See **Table 1** for details). We selected these targets because of their association with genes showing particularly high levels of noise (and thus, more likely to be driven by tuning) in single-cell proteomic analysis [30]. In our screening, homozygous replacements of HST3, HST4, SWR1, ISW2, and UBP8 with a kanMX cassette showed little effect on colony formation rates on ura-/6AU plates, and SIN3::kanMX/SIN3::kanMX strains showed severely compromised cell survival under growth-arrested conditions; all were excluded from further analysis. On the other hand, we found that genetic perturbations to the histone acetylation machinery through deletion of the key histone acetyltransferase GCN5 essentially abolished tuning, whereas deletion of the histone chaperone ASF1, in contrast, increased tuning rate by more than an order of magnitude (**Figure 6B**). At the same time, we show that the observed tuning process does not rely on transcriptional memory mechanisms grounded in chromatin localization, given the lack of effect of a NUP42 deletion (**Fig. 6B**; *cf.* [31]).

**Table 1:** Summary of genetic perturbations tested for effects on tuning rates.

| Perturbation | Direct effect | Effect on tuning |
| --- | --- | --- |
| GCN5::kanMX | Deletion of histone acetyltransferase subunit (acts in ADA, SAGA, SLIK/SALSA complexes) | **Inhibits** |
| SWR1::kanMX | Deletion of H2AZ exchange factor | No effect |
| UBP8::kanMX | Deletion of SAGA complex de-ubiquitinase | No effect |
| SIN3::kanMX | Deletion of Rpd3S/L histone deacetylase components | No effect |
| HST3::kanMX | Deletion of Sir2-family histone deacetylase | No effect |
| HST4::kanMX | Deletion of Sir2-family histone deacetylase | No effect |
| ISW2::kanMX | Deletion of DNA translocase involved in chromatin remodeling | No effect |
| ASF1::kanMX | Deletion of nucleosome assembly factor | **Accelerates** |
| NUP42::kanMX | Deletion of nuclear pore complex component known to be involved in transcriptional memory | No effect |

### Variations in colony formation rate are not a result of changes in viability

In interpreting our data on the effects of genetic perturbations on tuning (**Figure 6B**), it was crucial to consider the possibility that cells may lose viability at variable rates under different conditions, which could contribute to the observed differences in colony formation rates. We thus performed experiments to measure the rate of cell death in the presence of uracil starvation, and compared the results with the different colony formation rates observed. As shown in **Figure S5**, the effects of a mutation on survival and tuning rates are not significantly correlated. For example, deletion of GCN5 resulted in the nearly complete loss of transcriptional tuning, deletion of NUP42 had no effect, and deletion of ASF1 substantially enhanced tuning, yet none of these mutations shows a change in survival rates during incubation in uracil-free media compared with wild type cells sufficient to explain the observed change in colony formation rate (**Figure S5**). Even for the poorest surviving strain, GCN5::kanMX/GCN5::kanMX cells, colony formation rates after ten days are 100--1000 times lower than wild type cells even though survival rates are lower only by a factor of ten.

### Chemical perturbation of histone deacetylases inhibits the maintenance of the tuned state

Given the apparent importance of chromatin modifications in fitness-directed tuning, we also tested the effects of nicotinamide treatment (which inhibits the sirtuin class of histone deacetylases, or HDACs [32]) on reversion of the tuned cells back to a naïve state. As shown in **Figure 6A**, we found that chemical inhibition of sirtuin HDACs by nicotinamide treatment substantially accelerated reversion of tuned cells to the naïve state, further highlighting the importance of histone modification in transcriptional tuning. Our results demonstrate a crucial role for chromatin modifications in the establishment and maintenance of the tuning process, although the molecular details cannot be discerned from these data alone.

## Discussion

We have described a mechanism of adaptation through fitness-directed optimization of gene expression. In numerical simulations, the proposed framework has the remarkable capacity to simultaneously tune the expression of thousands of genes, enabling optimization of fitness without directly sensing environmental parameters. The demonstration that a phenomenon consistent with fitness-directed transcriptional tuning operates in *S. cerevisiae* has important implications for the adaptation of eukaryotic microbes to novel or extreme environments where their genetically encoded regulatory networks become inadequate. However, we speculate that transcriptional tuning operates in parallel with conventional regulation even in frequently encountered environments. Indeed, hard-coded regulatory networks cannot have the encoding capacity to optimally respond to every conceivable subtle change in the environment—even within the native habitat. We, therefore, favor a model in which dedicated regulatory networks quickly move the system to a state reasonably well matched to a given condition, and transcriptional tuning subsequently optimizes expression to achieve precise adaptation in every individual encounter.

The proposed mechanism is reminiscent of the biased random walk phenomenon in bacterial chemotaxis, where stochastic transitions in the rotation of the flagellar motor are biased towards the direction that increases chemoattractant signaling over time [33]. Detailed molecular mechanisms of chemotaxis have been worked out over the course of the last few decades, demonstrating the versatility of molecular processes in implementing rather complex computations (reviewed in [34]). Although our main focus here has been on establishing the phenomenology of fitness-directed transcriptional tuning, we have already identified some critical components. In particular, histone acetylation/deacetylation (via GCN5 and sirtuins) seem to play a central role, as deletion of GCN5 almost entirely abolished tuning. This is consistent with the high degree of intrinsic noise exhibited by the genes that are regulated by the SAGA complex, in which GCN5 is the catalytic subunit [30]. Previous work has shown that increased transcriptional noise is beneficial for adaptation to acute environmental stress [16]. Interestingly, however, early work demonstrated that deletion of GCN5 further increases expression noise in the context of the PHO5 promoter [35].

Taken together, these data suggest that transcriptional tuning is not driven by noise alone; rather we support a model in which the proper integration of noise, transcriptional memory, chromatin modification, and cellular-health feedback work together to implement a directed search mechanism to drive the expression level of individual genes to levels that maximize the internal health of the cell. Indeed, histone modification is tightly coupled with gene expression. Co-transcriptional histone modification can store recent memory of transcriptional activity [19, 20] and histone modification can, in turn, affect transcription rate [36]. There has been a longstanding debate on the functional significance of this reciprocal coupling. Our model and results help to unify these phenomena and demonstrate their functional relevance as requisite components of a transcriptional tuning-based cellular adaptation framework.

We note that our experimental setup for demonstrating transcriptional tuning has superficial similarities to a series of experiments performed in *S. cerevisiae* by the Braun lab, in which they sought to determine whether glucose-driven repression of the GAL1 promoter could be overcome to allow expression of a HIS3 construct in glucose-containing media [37, 38]. While the authors observed consistent emergence of growth in a large fraction of cells that they initially noted could be attributed to either genetic or epigenetic mechanisms [38], subsequent analysis has shown that in that experimental system, genetic mutations are the primary mechanism of adaptation, possibly driven by hypermutability of the genes involved in the response of interest [39-41]. These mutational mechanisms stand in clear contrast to the rapidly reverting epigenetic transcriptional tuning observed in our experiments.

In addition to perception of environmental parameters, cells also possess a variety of hard-wired homeostatic mechanisms sensing and responding to internal parameters, optimizing resource allocation in response to parameters such as growth rate [42-46] and metabolite/nutrient pools [47, 48]. However, while these mechanisms allow cells to sense their internal state, they still reflect specific evolved responses to alter resource allocation and gene expression in a *predefined* way in response to stress, standing in contrast with the ability of the transcriptional tuning process that we have outlined to enable a form of innovation of new transcriptional profiles in response to new environmental conditions.

The widely varying tuning rates for different promoters (**Figs. 3B-C and S1**) clearly indicate that sequence features can influence tuning efficacy. By design, all but one promoter driving URA3 in our experiments contained a TATA box, which has been linked to high intrinsic noise [30], condition-specific expression variability [28] and reliance on chromatin-mediated regulation [49, 50]. Indeed, replacement of the (TATA-containing) P_SAM3_ derived sequence in synprom with a similarly generated sequence from the TATA-free P_ARF1_ promoter substantially reduced tuning rates under the conditions tested (**Figure S1**). Exploring the relevance of other features, such as propensity for nucleosome positioning, will be important in future work.

Fitness-directed transcriptional tuning requires feedback of the global state of health to every promoter in the genome. The dependence of many histone modification enzymes on metabolic intermediates and cofactors (e.g., NAD+ for the sirtuin family of histone deacetylases [51, 52]; SAM for histone methyltransferases [53], and acetyl-CoA for histone acetyltransferases [54]) provides support for potential direct feedback of global fitness-related parameters to the epigenome [55, 56], and indeed we showed that chemical manipulation of sirtuin activity had substantial effects on retention of epigenetic memory. These enzymes may very well serve as distinct channels of health-related information utilized by transcriptional tuning. Alternatively, cells may utilize a single global health integrator (such as the mTOR system) as hypothesized in our idealized model. The mTOR pathway integrates diverse parameters of internal health including energy, nutrient availability, and cellular stresses [22]. Intriguingly, the mTOR pathway has recently been shown to regulate histone acetylation states through a variety of mechanisms [57, 58]

Fitness-directed transcriptional tuning has important implications for gene regulation. Beyond a potentially widespread mechanism of cellular adaptation, transcriptional tuning brings together seemingly unrelated phenomena under a unifying conceptual framework. These are areas of study at the frontier of genetics and biochemistry, including stochastic gene expression, transcriptional memory, and metabolic modulation of epigenetic states. Transcriptional tuning likely initially evolved as a mechanism for adaptation of single-cell eukaryotes to extreme environments. However, once available, it may have found utility as a versatile mechanism for controlling and fine-tuning gene expression in the context of physiological and developmental processes in metazoans, explaining the evolutionary arc of an ancient set of molecular mechanisms that now serve as key mediators of differentiation [59-61]. Exploring this possibility represents an important area for future research. Optimization of cellular health through the fitness-directed transcriptional tuning mechanism may also play an important role in allowing cancer cells to survive and thrive in a variety of microenvironments unfamiliar to their evolved regulatory networks, and in the face of extreme challenges imposed by chemotherapy and radiation. Indeed, transcriptional tuning provides a ready explanation for the wealth of recent evidence showing epigenetically mediated metastatic potential and chemotherapy resistance in a variety of cancer types [62-67].

## Materials and Methods

### Media and Strains

For routine growth of strains, we used YPD broth (10 g/L yeast extract, 20 g/L peptone, 20 g/L dextrose) or YPD agar plates (YPD broth + 20 g/L Bacto agar). We used standard recipes based on SC+glucose (SC+glu) [68] for all physiological experiments. SC/loflo refers to SC made with low fluorescence yeast nitrogen base (US Biologicals). In the case of SC+glu, we used dropout supplement powders interchangeably from ForMedium (DSCK012) and US Biologicals (D9515), although they differ slightly in the concentrations of adenine and *para-*amino benzoic acid supplied. SC+glu derivatives lacking particular supplements are specified as SC+glu-NUTRIENT; e.g., SC+glu-ura for SC+glu lacking uracil. We also refer to the commonly used mixture of SC+glu-ura with 6-azauracil added as ura-/6AU*i*, where *i* is the final concentration of 6AU in microgram/mL. The agar for all plates used in physiological experiments was either Noble agar (Difco) or quadruple-washed Bacto agar. For the removal of the GAL-GIN11 cassette in counter-selections (see below), cells were plated on YPGA agar plates (10 g/L yeast extract, 20 g/L peptone, 20 g/L galactose, 20 g/L agar, 100 microgram/mL ampicillin). All growth was at 30° C; liquid phase growth included shaking at 200-220 rpm in an Innova 42 incubator (New Brunswick).

As diagrammed in **Fig. 3A**, we constructed two classes of insertion cassettes. Each follows the pattern of having a promoter, a functional reporter protein fused to a fluorescent protein, and then ends with a CYC1 terminator. For URA3, the native sequence from *S. cerevisiae* was used, with the exception of one silent SNP and an A160S mutation that does not appear to alter enzyme function. The red fluorescent protein mRuby is described in [25]. For DHFR, we used murine DHFR from pSV2-dhfr [69] with an L22R mutation making it methotrexate-resistant [70]. GFP refers in all cases to superfolder GFP [26] codon optimized for *S. cerevisiae* using web-based tools from IDT (Integrated DNA Technologies). In each case, the reporter and fluorescent protein were separated by a short A/G/S containing linker. All constructs were cloned in bacterial hosts using pBAD-derived plasmids; separate plasmids were constructed with each promoter of interest downstream of a region homologous to the upstream target site in the *S. cerevisiae* genome, and URA3-mRuby-cyc or DHFR-GFP-cyc upstream of a region homologous to the downstream target site in the *S. cerevisiae* genome. All constructs were chromosomally integrated at the leu2Δ0 locus of our yeast strains. Double-stranded DNA for transformation in yeast was then generated by first amplifying the promoter and reporter constructs separately, using primers yielding 20-40 bp overlaps; we then used crossover PCR to generate the complete construct of interest and subsequent amplification to generate a sufficient quantity for transformation. All PCR used for strain construction was performed using Q5 high fidelity polymerase (NEB); routine PCRs for strain validation were instead performed using OneTaq or Taq polymerase (NEB).

Promoters for ADH1, HSP12, and RGI1 were cloned from our wild type strain (BY4743 or its haploid progenitors BY4741/BY4742), and included the entire region from 1700-1800 bp upstream of the start codon to the base immediately prior to the start codon. The ADH1 promoter was selected as a classic constitutive promoter [71]; HSP12 and RGI1 were chosen as they show high variance in expression between conditions [27, 28], a characteristic thought to be favorable for transcriptional tuning. Synprom was designed in two stages: the bulk of the DNA is a 600 bp random sequence generated using a Markov model to match the trinucleotide frequencies present across all natural *S. cerevisiae* promoters. To this sequence we appended the 200 bp immediately prior to the start codon of SAM3, to provide native transcription and translation start sites. The resulting sequence was then modified to remove all recognizable binding sites for yeast transcription factors (TFs) as follows: we used the set of position weight matrices and match thresholds in ScerTF [72] to identify all recognizable TF binding sites in the promoter, and randomized the sequences of only those regions and their immediate surroundings until no recognizable TF binding sites remained. The resulting perturbed sequence is given as **Data S1**. The required sequences were synthesized as gBlocks from Integrated DNA Technologies and combined via Gibson assembly [73].

All yeast strains were derived from BY4741 or BY4742 [74], which includes a complete deletion of the URA3 ORF (BY4741: Mat **a** his3Δ1 leu2Δ0 met15Δ0 ura3Δ0; BY4742: Mat α his3Δ1 leu2Δ0 lys2Δ0 ura3Δ0). Insertions of URA3 or DHFR fusion proteins were always at the leu2Δ0 locus unless otherwise noted. To facilitate consistent insertion, we replaced the leu2Δ0 allele of BY4741/BY4742 with a LEU2-GAL-GIN11 cassette [75], which allows growth in leucine-free media but inhibits growth in the presence of galactose. We note that at least in our copy of the BY474x strains, the leu2Δ0 deletion runs only from ChrIII:84799—ChrIII:93305, rather than extending to position 93576 as annotated. Nevertheless, the deletion is sufficient to remove the entire leu2 ORF.

Strains containing the fusion proteins were constructed by transforming the LEU2-GAL-GIN11 containing cells with appropriate double-stranded oligos (see above) and selection on YPGA plates, allowing replacement of the LEU2-GAL-GIN11 cassette with the desired insert. Insertions were confirmed by PCR product sizing. Diploid strains were derived by mating one BY4741-derived (mat **a**) strain with one BY4742-derived (mat α), and subsequently plating on SC+glu-lys-met or SC+glu-lys-met-cys. All transformations were carried out using the LiAc-PEG-ssDNA method [76].

Knockout strains were generated by beginning from appropriate haploids containing either a leu2::promoter-URA3 or leu2::promoter-DHFR construct or simply leu2Δ0, amplifying an appropriate kanMX knockout cassette from the corresponding strain in the *S. cerevisiae* gene deletion collection [77], and selecting on YPD + G418 plates. We confirmed the presence of kanMX at the appropriate site and absence of the native gene by PCR. Diploid knockout strains containing appropriate deletions and a URA3-mRuby insertion at leu2Δ0 were generated by mating these haploids as noted above.

### Colony formation assays

Experiments showing colony formation rates over time all follow a common formula. Cells were grown overnight in SC+glu media, and then in the morning back-diluted 1:200 into fresh, prewarmed SC+glu. The cells were grown for four to five hours at 30° C with shaking and then pelleted, washed once with 25 mL deionized (DI) water, pelleted, washed with 1 mL water, pelleted, and resuspended in 1 mL water. Specified dilutions were made in DI water from this final cell suspension.

Cells were then either plated on full plates at pre-chosen dilutions (100 microliters of an appropriate cell suspension), or a dilution series was spotted onto appropriate agar plates (10 microliters per spot). Plates were imaged and counted every 1-2 days for the duration of the experiment (lasting between a few days and weeks, depending on the experiment in question). Plates were wrapped in parafilm after ∼3 days to minimize drying. Plating was performed identically on SC+glu plates (to establish the number of cells being plated) and plates containing one or more test conditions (e.g., ura-/6AU).

Cells were counted either directly from the plates or from stored digital images. Direct plate counts were done manually for all visible colonies; for those counted from saved images, we imposed a minimum size threshold of 0.2 mm in diameter (rounding up to the nearest pixel). Times for counts were rounded to the number of days since plating.

### Death rate assays

To determine the survival rates of cells undergoing uracil starvation in the presence of various other perturbations, we measured the death rates of cells lacking any copy of URA3 in SC-ura+glu media. Cells were pregrown and washed as described above for plating assays, but then resuspended in liquid SC-ura+glu media and incubated at 30° C. Aliquots were regularly removed and spotted on SC+glu plates to determine the number of viable colonies. Survival rates are for leu2Δ0 homozygotes (the original BY4743 diploid, possibly with a homozygous deletion of a specified gene) with no available copy of URA3.

### Flow cytometry

Cells were analyzed by flow cytometry on an LSR Fortessa (Becton Dickinson) at the Columbia University Microbiology and Immunology Flow Cytometry Core Facility or University of Michigan Flow Cytometry Core. Cells to be used in these experiments were initially prepared and washed following the same pregrowth procedure as given above for colony formation assays, except that growth was in low fluorescence SC/loflo media instead of SC. A flask containing 25 mL of prewarmed media (generally ura-/6AU5 made from an SC/loflo base) was then inoculated with 200 microliters of the cell suspension, and cells were grown with shaking at 30° C. Subsequent data acquisition varied depending on the experiment to be performed.

For the long time courses shown in **Figure 4** and **Figure S2**, for an initial timepoint, 200 microliters of the washed cell suspension were combined with 500 microliters of 2x PBS/E (1x PBS with 10 mM EDTA added), 290 microliters DI water, and 10 microliters of flow cytometry counting beads (Invitrogen CountBright beads). At subsequent timepoints, snapshots were taken by combining 490 microliters of the growing cells, 10 microliters counting beads, and 500 microliters 2x PBS/E. In either case, cells were run on the Fortessa, with signals recorded for forward and side scatter, mRuby (using the Texas Red laser/filter set), and GFP (using the FITC laser/filter set).

Data were analyzed using the flowCore and flowViz modules of R [78, 79]. Beads and cells were first identified based on their forward scatter and side scatter (FSC/SSC) values (using permissive gates that capture the vast majority of each population) and fluorescence (beads were required to show very high fluorescence). For each growth phase (exponential in SC+glu, starving in ura-/6AU, growing in ura-/6AU), we obtained empirical autofluorescence corrections by analyzing populations in a similar growth state lacking the fluorescent tag on URA3. Guided by exploratory analysis, we fit a linear model for starving cells predicting mRuby and GFP autofluorescence as a function of the observed forward and side scatter, and used constant autofluorescence values characteristic of each of the two growing phases (obtained from cells with no fluorescent protein in a similar physiological state, either uracil-starved or undergoing transcriptional tuning-driven growth). During analysis of liquid phase fluorescent populations (shown in **Figs. 4** and **S2**), the predicted autofluorescence values were subtracted from the observed value; in these cases an additional gate was applied to remove events with very low forward scatter values, which had a very high variance in fluorescence and were well below the size of the main population.

For the use of FACS followed by plating to test the colony formation rates of highly fluorescent cells, cells were prepared as described above, sorted using a BD FACSAria, and then subsequently plated in equal quantities on SC+glu and ura-/6AU15 plates.

For the short timescale tuning data shown in **Figure 5**, the cells were grown for 3-4 hours side by side in SC/loflo+glu and –ura/loflo/6AU1 media, and then placed on ice and run directly on the flow cytometer. For each biological replicate (performed on different days), we grew leu2::synprom-URA3-mRuby/leu2::synprom-DHFR-GFP and nonfluorescent leu2::URA3/leu2Δ0 cells in parallel to allow direct comparison of the observed fluorescence levels. Analysis was performed separately for each biological replicate. We first normalized all fluorescence signals by the FSC-A signal raised to the power of 1.5, which we found empirically to be an effective correction removing most of the dependence of the fluorescence on cell size. Next, a mapping of FSC signals to expected autofluorescence on each channel was fitted using the R loess function (with default parameters), and the expected autofluorescence subtracted from the observed value for each cell to yield what we refer to as the blanked fluorescence. We then calculated and compared the changes in the median blanked fluorescence of the populations for the same cells grown in SC+glu vs. ura-/6AU1 media. Confidence intervals were calculated by bootstrapping with 200 bootstrap replicates.

### Whole genome sequencing

Cells for whole genome sequencing were taken directly from the growth condition of interest (ura-/6AU15 plate or ura-/6AU5 liquid media) and flash frozen in 15% glycerol or 1x TES (10 mM Tris, pH 7.5; 10 mM EDTA, 0.5% SDS). One reference sample grown under unselective conditions was taken for each starting strain to use as a baseline. Genomic DNA was isolated using a YeaStar Genomic DNA kit (Zymo Research) according to the manufacturer’s instructions. Samples were then barcoded and prepared for sequencing using a Nextera XT kit (Illumina, Inc.) and sequenced as part of a pooled library on a NextSeq (Illumina, Inc.).

Sequencing reads were clipped to remove adapters and commonly observed artifactual end sequences with cutadapt [80], and then further trimmed using Trimmomatic 0.30 [81] to remove very low quality (<3) end bases, retain only the portion of the read with a quality score above 15 in a 4 base sliding average window, and remove reads less than 10 bp long. Surviving trimmed reads were then aligned to the reference genome using Bowtie 2.1 [82]; the reference genome was constructed from the *S. cerevisiae* S288c genome (GenBank BK006934 – BK006949), deleting the URA3 ORF and inserting the sequence for the appropriate URA3 and DHFR constructs in separate copies of chromosome III at the LEU2 locus.

After alignment, mutational calls and read depths were obtained using the mpileup and depth modules of samtools 0.1.18 [83], respectively. Reads for called variants within 25 kb of the insertion site were examined manually and compared to the sequenced parental strain; validated variants are listed in **Table S1**.

### RNA isolation

RNA was isolated using an adaptation of the hot acid phenol method [84]. Cells for RNA isolation were grown under appropriate conditions (either in liquid phase or on agar plates), and then snap-frozen in 1x TES (10 mM Tris, pH 7.5; 10 mM EDTA; 0.5% SDS) and stored below -70° C. Snapshots of 200 to 600 microliters were taken from growing liquid phase cultures, whereas from agar plates we harvested 1-20 colonies of <0.5 mm diameter taken from the same plate as each biological replicate. RNA was isolated by rapidly thawing the cell suspension and mixing 1:1 with a 5:1 acid phenol:chloroform solution, then incubating 60 minutes at 65° C with occasional vigorous vortexing. The solution was then chilled on ice for 5 minutes, and centrifuged 5 minutes at 16,000 x g at 4° C. The aqueous phase was mixed 1:1 with additional acid phenol:chloroform, chilled, and centrifuged as before. The aqueous phase was then mixed 1:1 with a 24:1 chloroform:isoamyl alcohol solution, and centrifuged 5 minutes at 4° C. The resulting aqueous phase was transferred to a fresh tube and combined with 1/10 volume 3 M sodium acetate, 2 volumes of 1:1 ethanol:isopropanol, and 1/800—1/200 volume Glycoblue (Ambion), and then precipitated for at least 1 hour at -20° C and then at least 1 hour at -80° C. RNA was recovered by centrifuging 15 minutes at 16,000 x g at 4° C, washed with ice cold 75% ethanol, spun an additional 5 minutes, and then air-dried and resuspended in RNAse-free water. The samples were then further purified using a Zymo RNA clean & concentrator 5 according to the manufacturer’s instructions, including an on-column DNase digestion.

### Quantitative RT-PCR

Total RNA was purified from cells in the desired growth condition using the hot acid-phenol procedure described above. cDNA pools were generated for each sample using random hexamer-primed reverse transcription with Protoscript II (New England Biolabs) following the manufacturer’s instructions. cDNA pools were used directly in qPCR reactions without further purifications, assembling reactions using iTaq Universal SYBR Green Supermix (BioRad) following the manufacturer’s instructions, in GeneMate PCR plates. Plates were sealed with Microseal ‘B’ adhesive film (BioRad) and run on a BioRad CFX96 detection system. C_t_ values calculated by the instrument software were then exported for subsequent analysis. All isolated RNA was quantified on a Bioanalyzer (Agilent) and found to have an RIN >= 6.8.

For comparison of URA3 and DHFR expression, we calculated separate ΔC_t_ values for each qPCR run replicate by taking the median of all technical replicates from that run. Values plotted in **Fig. S3** reflect ΔCt data from 1-2 technical replicate wells on each of two to four separate, independently assembled runs; we plot the median of day-wise data points for each separate biological sample. Primer locations and sequences are given in **Table S3**. We performed a no-reverse transcriptase control reaction for each sample to ensure that DNA contamination did not contribute to the observed signal (data not shown).

qRT-PCR data were analyzed using a Bayesian hierarchical model treating the ΔC_t_ value between the URA3 and DHFR primers as follows:

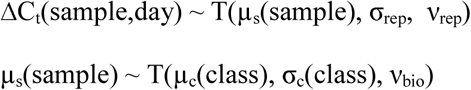

Parameters not otherwise specified were assigned appropriate uninformative priors. Here “sample” refers to a single biological sample and “class” to a single growth condition. The key parameter of interest is μ_c_ for each class of cells under study, the overall average URA3:DHFR difference for cells grown under that condition. We fitted the model using JAGS [85], and then report credible intervals and other inferences from the posterior distribution on μ_c_. Each of the ΔC_t_(sample,day) values used the median across 1-2 technical replicates for each primer pair.

### Cell count data analysis

Data were analyzed using custom-written python and R scripts. Uncertainties for cell counts (shown in plating and flow cytometry data) were calculated by treating each observed count as a Poisson random variable; using Bayesian inference with the Jeffreys prior [86], the posterior distribution for the rate parameter I (the concentration of cells) is given in closed form by

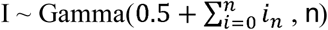

Where *n* is the number of observations and the *i*_*n*_are the observed counts. Error bars then indicate a central 95% credible interval for I given the observed data.

### Recovery experiments

Experiments to examine the reversion of tuned colonies toward a naïve state were performed as shown in **Figure S4**. Single colonies from a ura-/6AU15 plate were streaked out onto SC+glu and allowed to grow. From that plate, single colonies were again picked and underwent repeated passages in liquid media; each “passage” refers to a 200-fold dilution, which is then allowed to grow for 48 hours. Cells were also taken for plating from the original ura-/6AU15 plate, the first SC+glu plate stage, and several subsequent time points during liquid culture. Cells taken from plates were immediately diluted in water and spotted on SC+glu and ura-/6AU15 to track colony formation rates; cells taken from liquid passages were streaked out on SC+glu plates prior to use in spottings, in order to obtain a consistent physiological state. Plots for “naïve” cells refer to cells treated identically, except that they had initially been grown on SC+glu plates instead of ura-/6AU15 plates. Recovery was assessed based on the amount of time required for 1 in 10,000 cells spotted on the new ura-/6AU15 plate to form countable colonies (using linear interpolation of colony counts between observed data points); in the event that one dilution yielded no colonies passing our size threshold, but the next (10-fold more concentrated) spot gave an uncountable haze, we assigned a count of 1 to the more concentrated spot.

### Numerical simulations

The numerical simulations shown in **Figs. 2** and **5** were performed by implementing the model described in the text using the Matlab programming language and simulated using Matlab (Mathworks, Inc.) or GNU Octave version 3.8.1 [87], with qualitatively equivalent results obtained in either case. All simulations were performed using the same initial conditions (but different random seeds, for the sampling shown in **Fig. 5**).

## Author Contributions

Conceptualization, S.T.; Methodology, P.L.F. and S.T.; Software, P.L.F. and S.T.; Investigation, P.L.F., J.Y., and S.T.; Writing – Original Draft, P.L.F., J.Y., and S.T; Supervision, S.T.

## Acknowledgments

We thank members of the Tavazoie laboratory and Sohail Tavazoie for helpful discussions and feedback on the manuscript. PLF was supported by a K99 Award from NIGMS. JY was supported by the NIH MSTP program at Columbia University Medical School. ST was supported by the NIH Director’s Pioneer Award.

## Supplemental Figure Legends

**Figure S1**: *Stochastic colony formation rates for cells with URA3 driven by a variety of synthetic promoters.* The “synprom” discussed in the main text is synprom5-sam3. Data are taken from SC-ura+glu agar plates containing the listed 6AU concentration. An ‘x’ followed by a dashed line indicates the threshold of detection from the experiment, marked at the last data point prior to any colonies being observed.

**Figure S2:** *Promoter-specific transcriptional tuning of URA3 expression by native promoters in S. cerevisiae.* Flow cytometry data on counts and fluorescence distributions for cells containing URA3-mRuby under control of the noted promoter (P_RGI1_ or P_HSP12_) and DHFR-GFP under control of P_ADH1_. Cells were grown in liquid ura-/6AU5 media. For the cell counts (top), colors correspond to different biological replicates started on different days. Arrows indicate two timepoints from each strain for which fluorescence cumulative distribution functions (CDFs) are shown below; in each CDF a given time-point (solid line) is compared to the distribution present for cells in logarithmic growth in SC+glu media (dashed lines). The values shown are log_2_ ratios to the median value of cells growing exponentially in SC+glu. GFP signals are shown in green and mRuby signals in red. Error bars for cell counts show central 95% credible intervals.

**Figure S3**: *Local tuning of URA3 expression.* RT-qPCR based quantification of URA3:DHFR ratio for synprom-URA3-mRuby/synprom-DHFR-GFP cells either in in SC+glu media (liquid or plates), from tuned liquid cultures in ura-/6AU5 media, or for tuned colonies on ura-/6AU15 plates. Large points show estimated averages obtained via a Bayesian hierarchical model (see Methods), small points show values for biological replicates, and error bars indicate 75% (strong) or 95% (thin) credible intervals. P(diff) in each case gives the posterior probability that the –URA value is greater than the corresponding SC+glu value. See **Table S3** for primer sequences.

**Figure S4**: *Recovery of cells taken from ura-/6AU plates toward a naïve state.* Experimental setup for assessing recovery to a naïve state. Tuned colonies growing on a –ura/6AU15 plate were streaked onto an SC+glu plate, colony purified, and then grown for 48 hour cycles in liquid SC+glu. Colony formation rates were assessed after each transfer, and quantified according to the amount of time needed for formation of one colony per 10,000 cells plated (see Materials and Methods for details).

**Figure S5:** *Survival of cells in –ura media in the presence of genetic perturbations*. Loss of viability of cells with the indicated gene deletion, but not bearing any copy of URA3, in SC-ura+glu media. Separate traces indicate different biological replicates.

